# Centralizing content and distributing labor: a community model for curating the very long tail of microbial genomes

**DOI:** 10.1101/031286

**Authors:** Tim Putman, Sebastian Burgstaller, Andra Waagmeester, Chunlei Wu, Andrew I. Su, Benjamin M. Good

**Affiliations:** Department of Molecular and Experimental Medicine, The Scripps Research Institute, La Jolla, USA; Department of Molecular and Experimental Medicine, The Scripps Research Institute, Micelio, Antwerp, Belgium

## Abstract

The last 20 years of advancement in DNA sequencing technologies have led to the sequencing of thousands of microbial genomes, creating mountains of genetic data. While our efficiency in generating the data improves almost daily, applying meaningful relationships between the taxonomic and genetic entities requires a new approach. Currently, the knowledge is distributed across a fragmented landscape of resources from government-funded institutions such as NCBI and Uniprot to topic-focused databases like the ODB3 database of prokaryotic operons, to the supplemental table of a primary publication. A major drawback to large scale, expert curated databases is the expense of maintaining and extending them over time. No entity apart from a major institution with stable long-term funding can consider this, and their scope is limited considering the magnitude of microbial data being generated daily. Wikidata is an, openly editable, semantic web compatible framework for knowledge representation. It’s a project of the Wikimedia Foundation and offers knowledge integration capabilities ideally suited to the challenge of representing the exploding body of information about microbial genomics. We are developing a microbial specific data model, based on Wikidata’s semantic web compatibility, that represents bacterial species, strains and the gene and gene products that define them. Currently, we have loaded 1736 gene items and 1741 protein items for two strains of the human pathogenic bacteria *Chlamydia trachomatis* and used this subset of data as an example of the empowering utility of this model. In our next phase of development we will expand by adding another 118 bacterial genomes and their gene and gene products, totaling over ~900,000 additional entities. This aggregation of knowledge will be a platform for community-driven collaboration, allowing the networking of microbial genetic data through the sharing of knowledge by both the data and domain expert.

## Introduction

The relatively small and non-repetitive nature of microbial genomes, coupled with the rapid advancement of sequencing technology in the last decade, have led to the generation of a staggering amount of bacterial genome records. The National Center for Biotechnology Information (NCBI) Genome Database currently maintains genome records for over ~3000 high quality reference and representative genome assemblies and another ~50,000 incomplete assemblies. The existing collections of genomes are just the beginning; The Earth Microbiome Project (http://www.earthmicrobiome.org) is in the early stages of analyzing and cataloguing over ~200,000 environmental samples from around the world, and estimates that this will result in the sequencing of ~500,000 reconstructed microbial genomes (1). Making sense out of this abundance of data, while a daunting challenge, will generate a wealth of knowledge for the microbial and human genomic research community.

For microbial genomes, as well as most other biological data, knowledge is distributed across resources that occupy the full spectrum from very large, broad coverage, centralized, major government-funded institutions such as NCBI and UniProt to boutique, topic-focused databases like the ODB3 database of prokaryotic operons, to the unstructured primary literature. The ability to smoothly process data from across that spectrum would greatly increase the efficiency of microbial research.

An example of such a question might be, “what other microorganisms influence the persistence of an infection by a human pathogen such as *Chlamydia,* and by what mechanism?”. An expert may generate hypothetical answers to this question by blending their knowledge with information spread through the literature and various databases. As an example, the statements illustrated in Figure 1, originating from multiple sources, including primary literature (2–5), and structured databases (NCBI Gene, UniProt, Drugbank, BRENDA), link together to yield the hypothesis that co-infection by *Prevotella spp., Clostridiales spp and Escherichia coli* in the vaginal microbiome increase the persistence of infection through the generation of indole(6,7), a key substrate in the Tryptophan biosynthesis pathway. Experts have done this leg work and generated the hypothesis that there is a greater risk of clearance failure, leading to persistent infection that should be treated appropriately, when these other indole-creating microbes are present (5).

**Figure 1.**
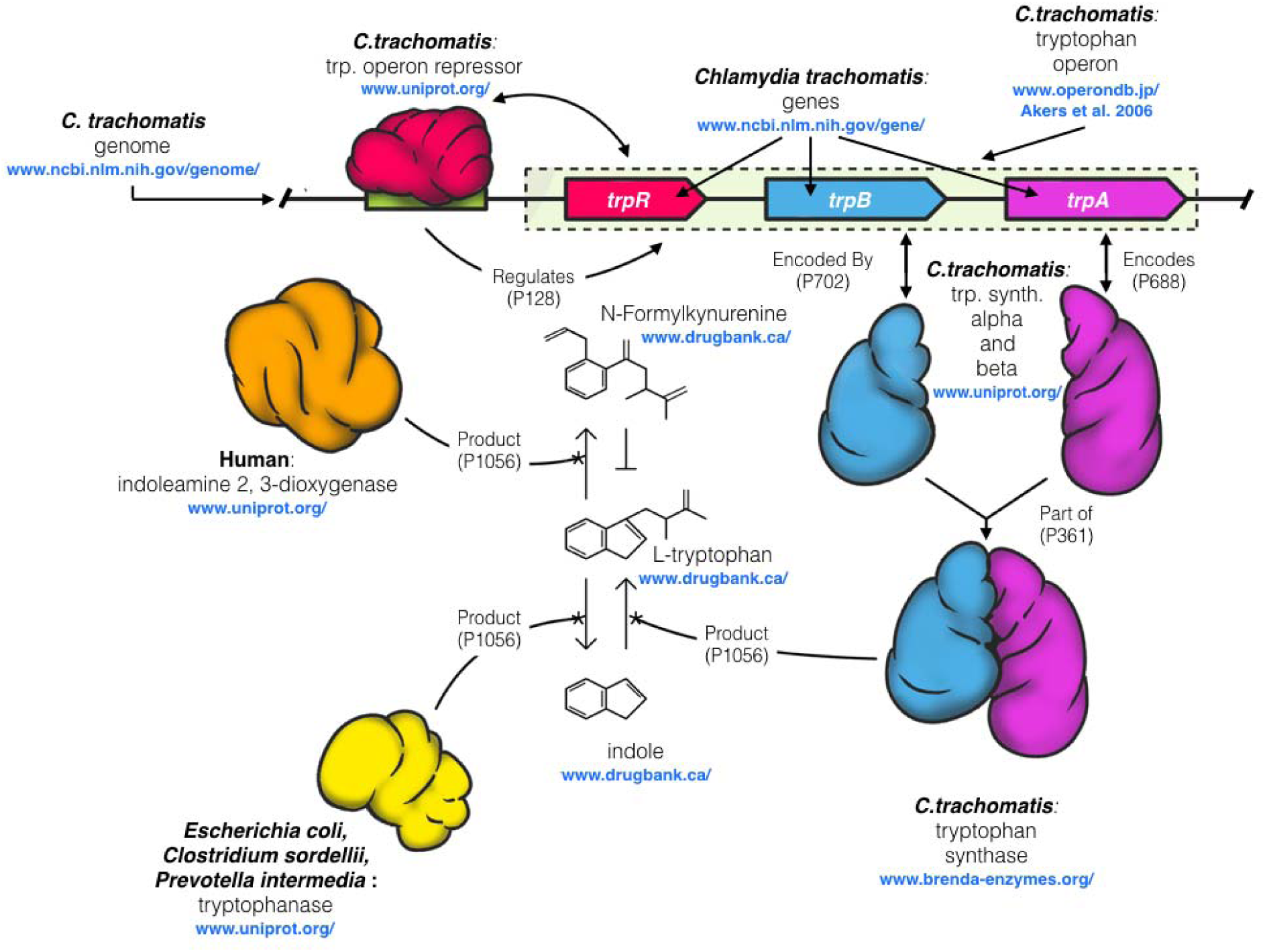
Illustration of the complex network of interacting entities between human, chlamydial, and other microbial species in the urogenital microbiome. When a human epithelial cell is infected by *C. trachomatis,* it responds by depleting the cell of L-tryptophan, an essential amino acid for chlamydial growth, through IFN-*γ* mediated expression of the tryptophan degrading enzyme indoleamine 2,3-dioxygenase (IDO)(orange) (2,3). IDO degrades tryptophan to N-Formylkynurenine, a tryptophan precursor that *C. trachomatis* is not capable of converting into tryptophan. Often this clears the infection, but episodically *C. trachomatis* rescues itself from this host defense by converting exogenous indole into L-tryptophan through gene expression regulated by its *trp operon* (*5*). Several experiments support the hypothesis that the likely source of exogenous indole is from other microbes in a perturbed vaginal microbiome; as part of L-tryptophan degradation via the pyruvate pathway. Microbes producing tryptophanase (yellow), an enzyme that degrades L-tryptophan to indole and pyruvate are commonly found in the urinary tract of patients also presenting with bacterial vaginosis (BV) (4). Example indole producers, commonly associated with BV in the female urogenital tract include *Prevotella spp., Escherichia coli, and Clostridiales spp.* (6–8). Blue URLs indicate the various resources that maintain the data. The arrows between entities indicate the properties used to define their relationships once aggregated in Wikidata.

By pulling these pieces of knowledge together into a common database, with defined connections between them, a list of the taxa involved can be generated as candidate answers to the above question with a single query. Once this is achieved and new data is added, the network grows and the collective benefit grows as well. The complicated and disordered is given order in a central container with a mechanism for sifting through it, giving the chlamydial researcher a powerful tool for making sense of the published data.

Model organism databases such as the Mouse Genome Database(http://nar.oxfordjournals.org/content/43/D1/D726.short) MGI, would greatly aid researcher’s ability to unlock connections between microbes and the organisms they interact with. However, large data warehouses such as this are typically maintained by expensive teams of data and domain experts. The immense scale of microbial data is economically incompatible with this kind of centrally-funded approach, and the same resolution would not be achieved. One way or another, the greater scientific community, encompassing both active scientists and interested members of the general public must be empowered to contribute their mental energy in a community-wide collaborative effort (9). Here, we propose that Wikidata may provide the means to achieve this goal.

Wikidata is a new, centralized, yet openly editable platform for semantic knowledge representation that is maintained by the Wikimedia Foundation (the same entity that maintains all of 200+ different language Wikipedias). Centralizing structured knowledge in this open database generates the opportunity to distribute the labor of data curation across a far broader community than was before realistic. In doing so, it offers a new approach to the knowledge integration problem that is ideally suited to the challenge of representing the exploding body of information about microbial genomics. Here, we describe the initial work of building a Wikidata-based representation of microbial genetics.

### Wikidata as a centralized microbial database

A centralized resource for microbial genomics will need to capture a wide variety of different kinds of entities and relationships to support useful queries. Rather than attempt to build a system that models all of this complexity up-front, we are taking the approach of seeding the openly extensible wikidata database with the beginnings of this model and thus encouraging the broader community to see the opportunity to collaborate on its evolution. Wikidata provides an ideal technical and social platform for undertaking this project. Its schema-free nature naturally supports data model changes and its open, wiki-based nature supports constructs such as ‘talk pages’, ‘watchlists’, and ‘wikiprojects’ that have proven effective in facilitating the attainment of community consensus over time in other open projects such as the GeneWiki (10).

As a starting point for seeding the collaborative creation of a centralized microbial database in Wikidata, we established the structures needed to represent the entities and relations depicted in Figure 1. In the context of Wikidata, this work amounts to the creation of a set of ‘items’ and ‘properties’ that are used to describe features of those items.

A Wikidata item is defined by a unique identifier (e.g. Q131065), a label (i.e. *Chlamydia trachomatis*), a description (i.e. ‘species of prokaryote’), and a set of ‘claims’ about the item organized into ‘statements’ (Figure 2). A statement consists of a triple with an item as the subject, a wikidata-defined property as the predicate (i.e. taxon rank, property P105), and another wikidata item or Literal data value as the object. Optionally, a set of references can be added as evidence and provenance for the claim made by the triple, and qualifiers can specify the context where the claim is valid (https://www.mediawiki.org/wiki/Wikibase/DataModel/Primer).

**Figure 2.**
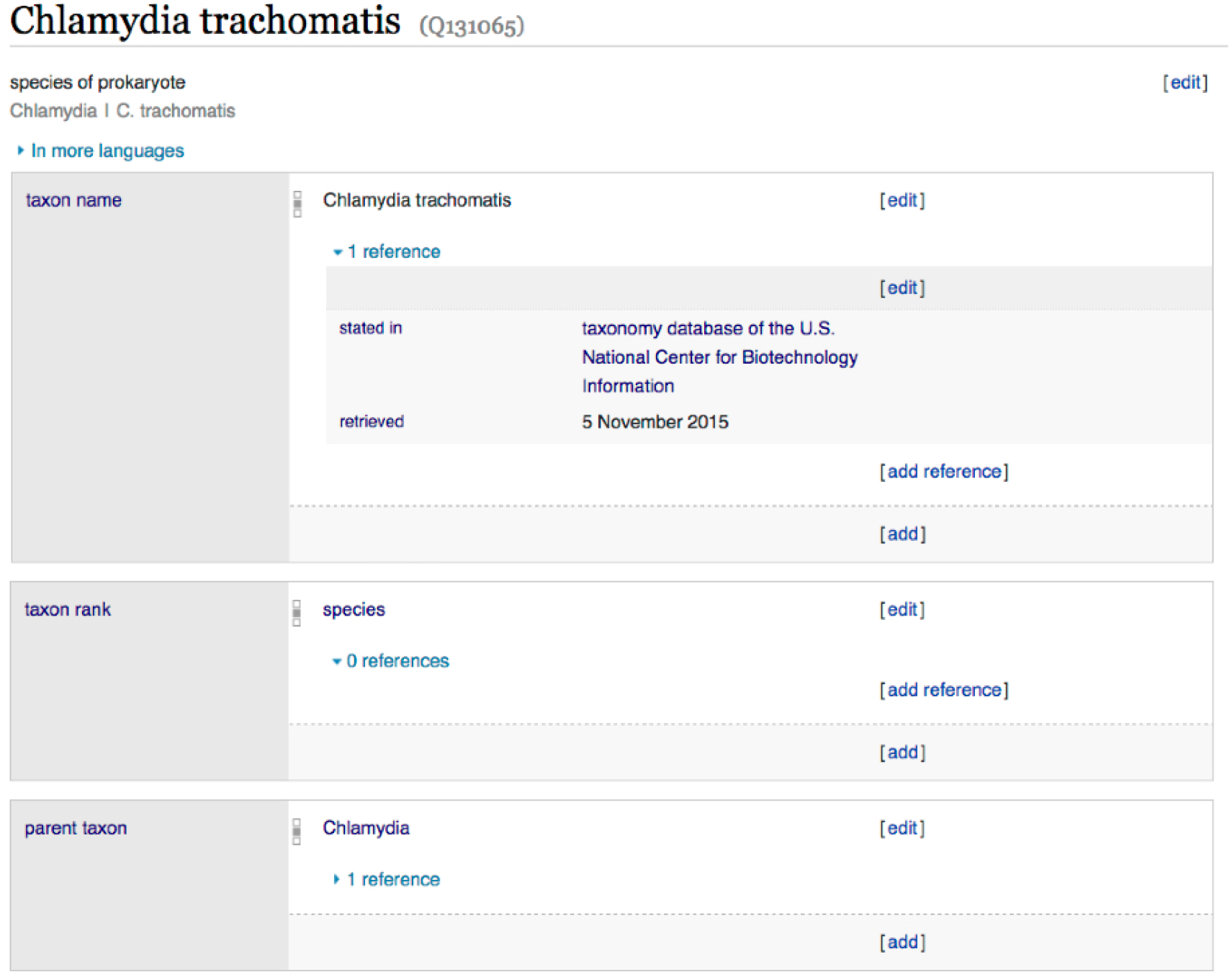
Example Wikidata item. A Wikidata item defined by its label, description, and statements that provide annotations and create relationships with other items in the database. (https://www.wikidata.org/wiki/Q131065).

The ‘ontology’ of Wikidata is determined by the set of properties that may be used to create claims about the items within it. Entities can be created at any time, but properties can only be created by elected administrators blessed with the privilege. When a new property is required, it must first be proposed and subjected to discussion with the community. Once consensus is achieved, the property is created by the administrator and is immediately available for use.

The properties needed to support our current data model are listed in Table 1. It is worth noting that most of these properties are either generic (i.e. subclass of) or defined by the Molecular Biology WikiProject (https://www.wikidata.org/wiki/Wikidata:WikiProject_Molecular_biology) prior to outset of the present work on microbial genomes. Already, wikidata is showing how an open system can evolve over time with subsequent efforts building directly on prior work.

**Table 1.** Wikidata properties in the microbial data model

| ID | Name | Value type |
| --- | --- | --- |
| P685 | NCBI Taxonomy ID | String |
| P105 | Taxon Rank | Wikidata Item |
| P171 | Parent Taxon | Wikidata Item |
| P2249 | RefSeq Genome ID | String |
| P1542 | Cause of | Wikidata Item |
| P351 | Entrez Gene ID | String |
| P279 | Subclass of | Wikidata Item |
| P703 | Found in taxon | Wikidata Item |
| P644 | Genomic Start | String |
| P645 | Genomic End | String |
| P702 | Uniprot ID | String |
| P637 | RefSeq Protein ID | String |
| P702 | Encoded by | Wikidata Item |
| P688 | Encodes | Wikidata Item |
| P680 | Molecular Function | Wikidata Item |
| P681 | Cellular Component | Wikidata Item |
| P682 | Biological Process | Wikidata Item |
| P361 | Part of | Wikidata Item |
| P128 | Regulates | Wikidata Item |
| P1056 | Product | Wikidata Item |

Some of the general purpose properties such as ‘product’, ‘part of’, ‘cause of’, and ‘regulates’ are currently used to establish the connections in Figure 1 but are likely to replaced or extended with more biology-specific relations (such as ‘precursor’ and ‘substrate of’) over time. The other aspects of the current model that are more specific to representing microbial data are depicted in Figure 3.

**Figure 3.**
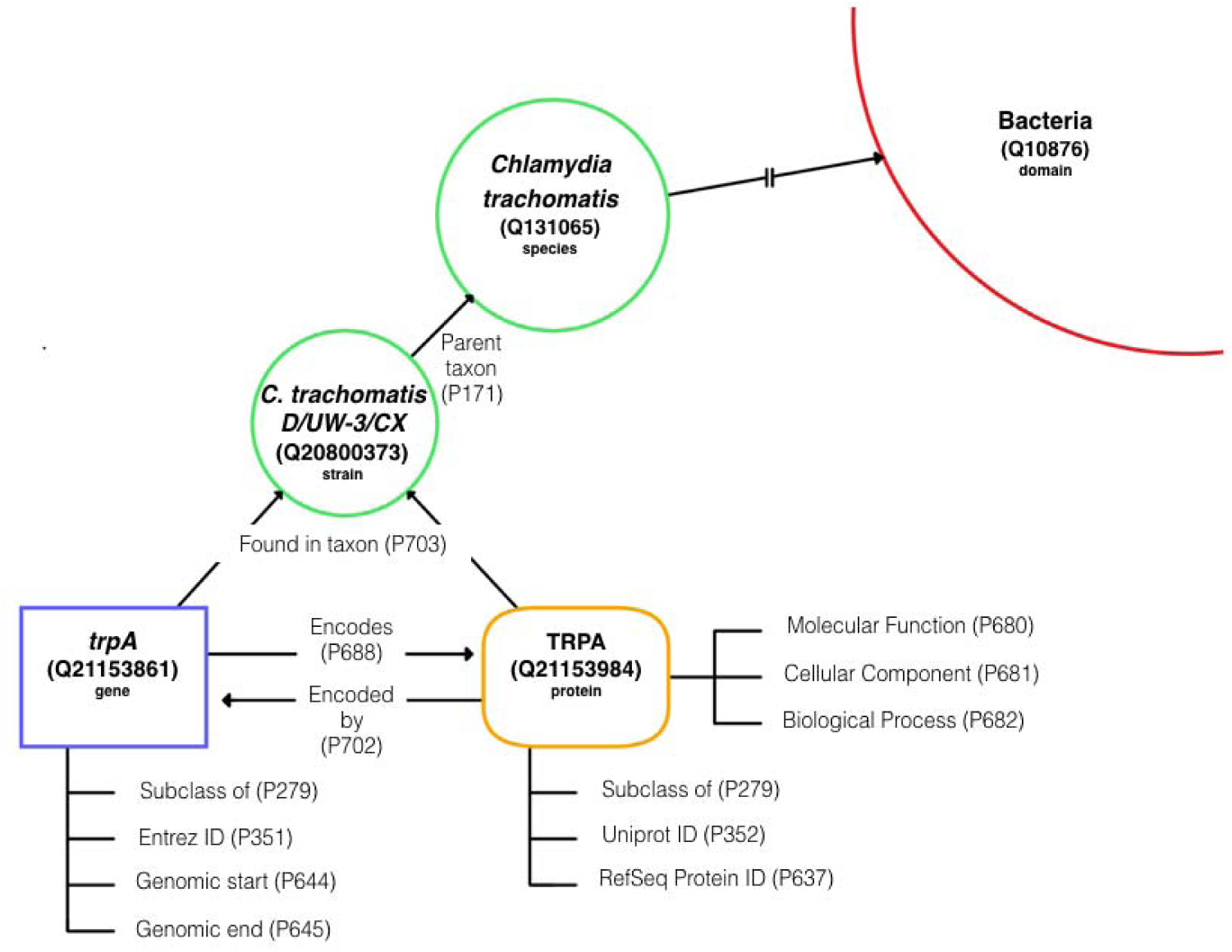
Data model template. The basic framework of the microbial genetic data model in Wikidata showing items and the statements that connect them. Item types are demarcated by label and color (i.e. gene item = blue and protein item = orange).

One key requirement for modeling microbial data is the capacity to represent multi-species, multi-strain datasets. Microbiome and genomic research require the ability to do both intra- and interspecies comparative analysis. To support this work, our model follows a hierarchical taxonomy ranking scheme with the microbial species assigned to a Wikidata item (i.e. *Chlamydia trachomatis* #Q131065) defined by the core properties ‘NCBI Taxonomy ID’ (P685) (‘813’), ‘Taxon Rank’ (P105)(‘species’) and ‘Parent Taxon’ (*P171*)(*‘Chlamydia*)

Since genome annotations are based on the genome assembly of the specific strain sequenced and that genome assembly has its own unique identifier (i.e. NCBI RefSeq Genome Accession number), strain level distinction is critical in bacteria. Individual strain items (e.g. Chlamydia trachomatis D/UW-3/CX #Q20800373) include the same core properties as a species item, the ‘RefSeq Genome ID’ (P2249), and are linked to the species item via the ‘Parent taxon’ (P171) property.

On the molecular level, the gene and protein must be kept as distinct entities, while maintaining their connections for queries down the line. A microbial gene item contains the similar core properties of a human gene item, including ‘Entrez Gene ID’ (P351) and ‘Subclass of’ (P279), but, ‘Found in taxon’ (P703) was added to distinguish which strain/genome assembly this particular gene came from. The gene links to its product item via the ‘Encodes’ (P688) property and reciprocally, the protein item will link to the gene that encoded it by the ‘Encoded by’ (P702) property. Core properties for microbial protein include ‘RefSeq Protein ID (P637), ‘UniProt ID’ (P352), ‘Found in taxon’(P703) and ‘Subclass of’ (P279). Functional annotations are downloaded from the UniProt protein record and included here as subclasses of the gene ontology terms, ‘molecular function’(P680), ‘cellular component’ (P681) and ‘biological process’ (P682)’.

### Populating and querying microbial genes in Wikidata

Given the data model depicted in Figure 3 and encapsulated in the properties listed in Table 1, we have seeded Wikidata with representative content for two chlamydial genomes (totaling 1736 gene items and 1741 protein items), from various public databases. This work was carried out with a ‘bot’, a program for making automated edits in Wikidata, with source code available at (www.bitbucket.org/sulab/wikidatabots/src) In addition, we manually established all of the Wikidata items and relationships needed to realize the operon data structure in Figure 1. This information can be accessed through the various APIs offered by Wikidata (https://www.wikidata.org/w/api.php, https://query.wikidata.org/). As an example, a user can easily retrieve all genes, proteins, and gene ontology annotations for the two strains of Chlamydia that are currently loaded using a wikidata SPARQL query (Figure 4).

**Figure 4:**
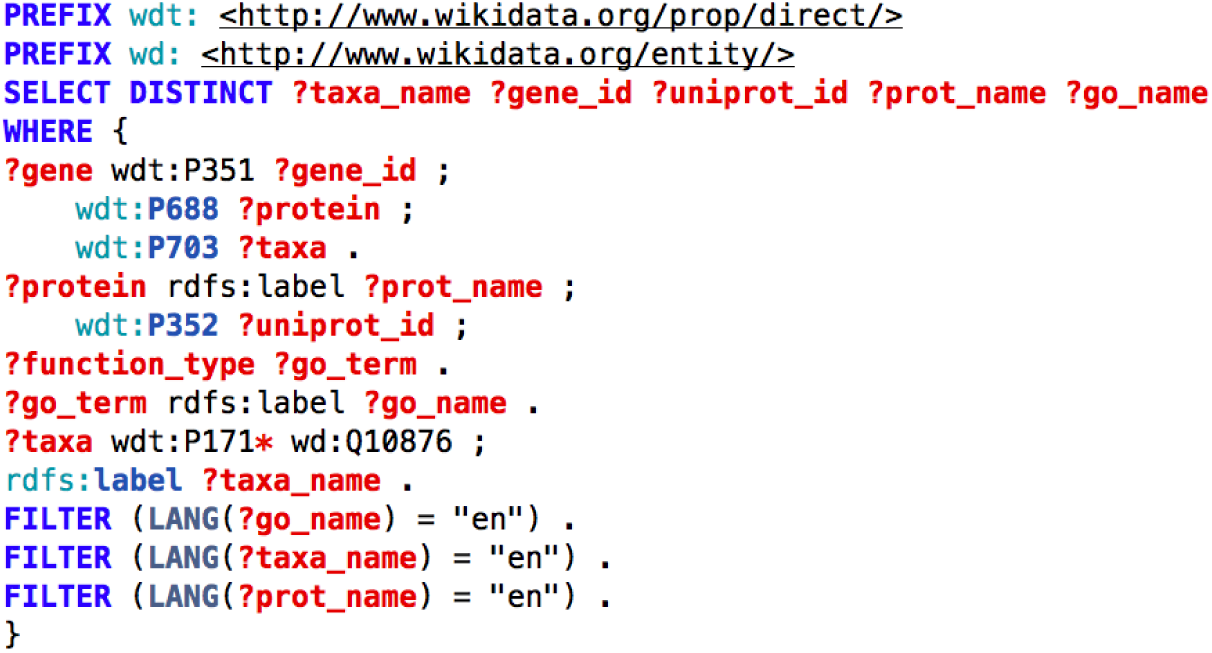
SPARQL query for all microbial genes, proteins and associated Gene Ontology annotations in Wikidata. Properties used: P351 = entrez_gene_id, P688 = encodes, P703 = found in taxon, P352 = uniprot_id, P171 = parent taxon. Note that the * operator on P171* results in a recursive search for organisms that descend from wd:Q10876 (Bacteria). This query may be executed at https://query.wikidata.org/.

Note that the query actually requests this information for all bacteria through the “?taxa wdt:P171* (parent taxons) wd:Q10876 (Bacteria)“ aspect of the query. As more bacterial genomes are loaded by us or other groups, the same query will return more and more data. As another example, the following SPARQL query returns all operons, their regulators, and their products (Figure 5).

**Figure 5:**
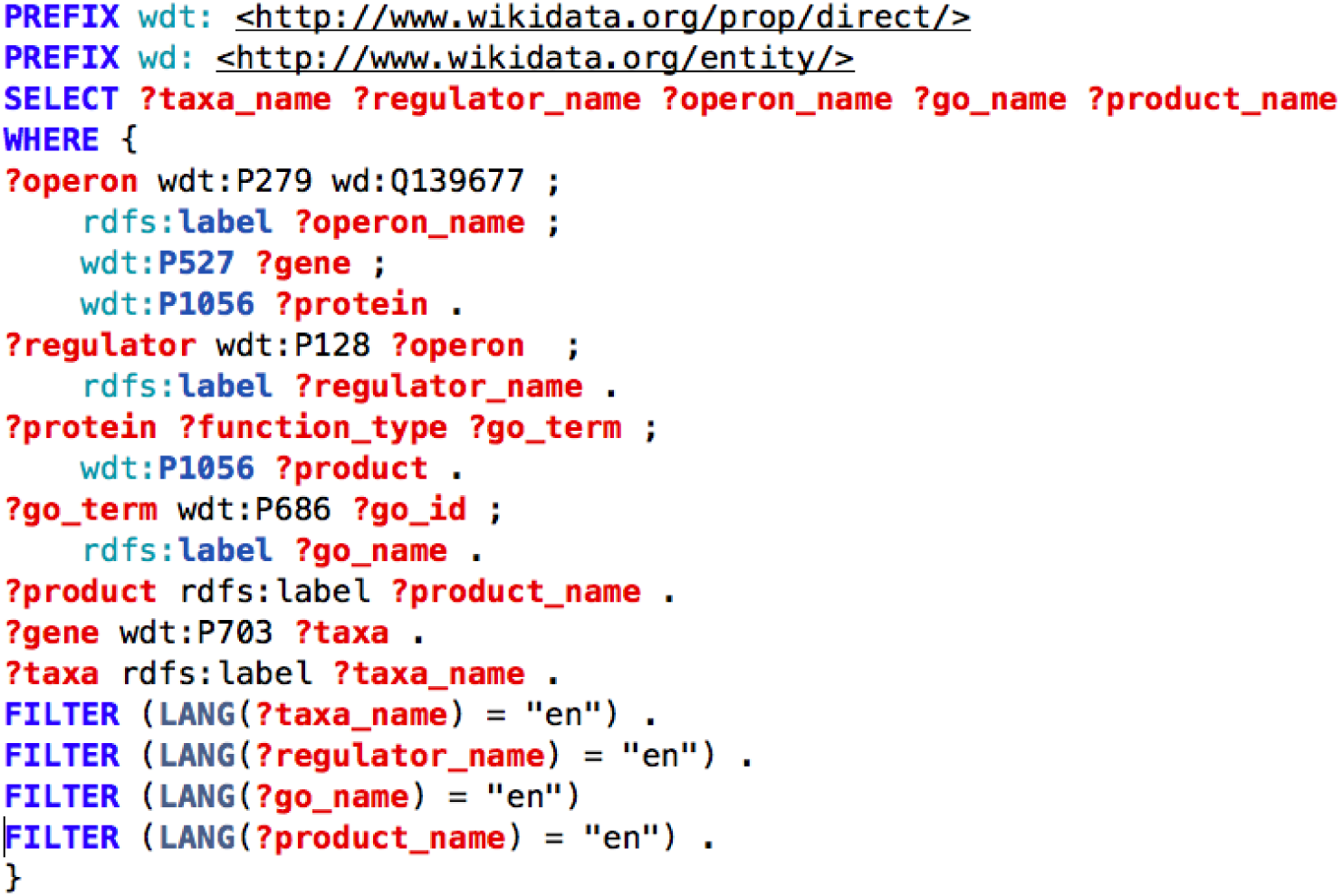
SPARQL query for all operons, their regulators, the taxon that expresses them and their functional products in Wikidata. Q139677 is the Wikdata item for the class ‘operon’. Properties used: P279 = subclass of, P527 = has part, P1056 = product, P128 = regulates, P688 = encodes, P703 = found in taxon. This query may be executed at https://query.wikidata.org/.

Revisiting the example question regarding organisms that are likely to be related to the persistence of chlamydial infections, we can ask what microbes are located in the female urogential tract and capable of generating indole as follows (Figure 6).

**Figure 6:**
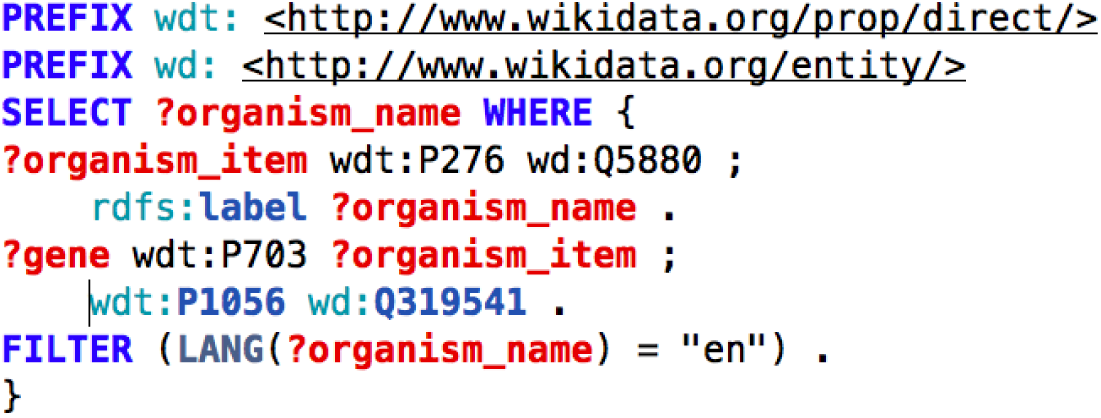
SPARQL query for all organisms that are located (P276) in the female urogential tract (wd:Q5880) and that have a gene with product (P1056) indole (wd:Q319541). This query may be executed at https://query.wikidata.org/.

## Discussion

Wikidata is certainly not a replacement for core data curation centers such as NCBI and UniProt. But it could form the basis for a complementary, stable, and cost effective approach for capturing content that is either left trapped in the literature or represented only in small databases subject to the perils of funding cuts and general link rot. Though the microbial queries listed above currently return only a small fraction of the relevant content that exists in the world, the power of the Wikidata approach is that our seedling database can be extended by anyone with the will to do so.

Wikidata is now edited by more than 15,000 active users and currently has over 15 million content pages (https://www.wikidata.org/wiki/Special:Statistics). Because of its open structure, its change tracking features, and its evidence-capturing data model, it encourages community participation at all levels. While the community consensus building process can be slow and at times frustrating, it drives the stability and quality of Wikidata content.

Topic-specific databases can lose funding and disappear (11). Even government backed institutions like NCBI and EBI are vulnerable to funding cuts depending on the political climate. The unique connection between Wikidata and all the Wikipedias already make it one of the most well-known and easily discoverable knowledge bases in the world (12,13). Data deposited here is far less likely to be lost, especially when care is taken to weave it into what already exists. Every item loaded to Wikidata (MediaWiki Foundation’s third most active project) becomes a fixed point in a stable, self-sustaining knowledge representation platform that anyone can add to, and anyone can help the network grow through sharing the benefits of their own expertise.

The open access, community driven nature, of Wikidata contributes to its perpetuity, but the major enduring factor is its universal utility. Wikidata is a place for knowledge of any conceivable topic from surfing (Q159992) to bacteria (Q10876). This variety of topics generates community support that a topic-specific, funding dependent database can not compete with. In addition to support, Wikidata creates the ability to link a surfboard (Q457689) to surfing (Q159992), the ‘sport’ (P641) it is used in, or *Chlamydia trachomatis* D/UW-3/CX (Q20800373) to pelvic inflammatory disease (Q558070), a disease it is the ‘cause of’ (P1542) in humans (Q5). Moreover, it in principle allows microbial genetics data to be linked to data from related fields, including pharmacology and epidemiology.

These relationship examples highlight another powerful virtue of Wikidata compared to other data storage platforms; adding data to Wikidata requires the use of meaningful properties for relating entities. It is insufficient to simple state “rdf:seeAlso” as the link between two related entities (as many major databases do in their RDF representations). A relationship between items can not be added without an appropriate property in place, requiring the data model to be defined prior to importing the data. This process of creating properties through community discussion and consensus drives the development of their ontology up-front, rather than forcing the burden of integrating ambiguous content downstream to consumers.

While Wikidata provides an excellent framework for housing some forms of data, it is not without its limitations. Not all content is appropriate for Wikidata. It is a database of referenced claims about the world and should not, for example, be a repository for sequences or expression data. There is no built-in reasoning in Wikidata. Editors cannot be constrained from making claims that may break data models spread across multiple items. As an openly editable resource, it is possible for data to be disrupted by edits from both well-intentioned editors and, at least theoretically, by malicious users (though true vandalism has thus far not happened at detectable levels).

Even in consideration of these limits, Wikidata is a tremendous potential platform for managing the process of collaboratively understanding microbial genomics. In support of this objective, our immediate next steps are to load the remaining 118 microbial reference genomes from NCBI (http://www.ncbi.nlm.nih.gov/genome/browse/reference/), encompassing bacteria that are the most studied and relevant to human health. This will load an additional ~900,000 entities between genes and gene products. These items will form a foundation upon which we invite the microbial research community to collaboratively synthesize their knowledge as it evolves into the future.

## Conclusion

We invite and encourage the rest of the scientific community to join our cause in creating this universal microbial genomics resource. We have shown that the aggregation of a subset of data enables powerful queries that demonstrate the potential of connecting data from fragmented sources into a centralized well-defined structure. In addition to data that can be collected from structured data sources and aggregated in Wikidata by bot, a great deal of important information resides only in primary literature. Accessing and integrating this content requires human editors. It is thus imperative that we engage the microbial research community to help build Wikidata to its potential; a centralized, semantic web compatible mechanism for making sense of mountains of microbial genetic data

## Acknowledgements

We would like to thank Lynn Schriml of the University of Maryland Baltimore, Baltimore, MD, United States for suggestions of resources for microbial data.

## Funding

This work is supported by the National Institutes of Health under grant GM089820, GM083924, GM114833 and DA036134.

